# How Healthy Older Adults Enact Lateral Maneuvers While Walking

**DOI:** 10.1101/2023.02.24.529927

**Authors:** David M. Desmet, Meghan E. Kazanski, Joseph P. Cusumano, Jonathan B. Dingwell

## Abstract

**Background:** Walking requires frequent maneuvers to navigate changing environments with shifting goals. Humans accomplish maneuvers and simultaneously maintain balance primarily by modulating their foot placement, but a direct trade-off between these two objectives has been proposed. As older adults rely more on foot placement to maintain lateral balance, they may be less able to adequately adapt stepping to perform lateral maneuvers.

**Research Question:** How do older adults adapt stepping to enact lateral lane-change maneuvers, and how do physical and perceived ability influence their task performance?

**Methods:** Twenty young (21.7 ± 2.6 yrs) and 18 older (71.6 ± 6.0 yrs) adults walked on a motorized treadmill in a virtual environment. Following an audible and visual cue, participants switched between two parallel paths, centered 0.6m apart, to continue walking on their new path. We quantified when participants initiated the maneuver following the cue, as well as their step width, lateral position, and stepping variability ellipses at each maneuver step.

**Results:** Young and older adults did not differ in when they initiated the maneuver, but participants with lower perceived ability took longer to do so. Young and older adults also did not exhibit differences in step width or lateral positions at any maneuver step, but participants with greater physical ability reached their new path faster. While only older adults exhibited stepping adaptations prior to initiating the maneuver, both groups traded-off stability for maneuverability to enact the lateral maneuver.

**Significance:** Physical and perceived balance ability, rather than age *per se*, differentially influenced maneuver task performance. Humans must make decisions related to the task of walking itself and do so based on both physical and perceived factors. Understanding and targeting these interactions may help improve walking performance among older adults.

## 1. Introduction

Falls are the leading cause of fatal and non-fatal injuries among older adults [1]. Falls to the side are particularly dangerous given both the significant risks posed by these falls [2] and that humans are thought to be more unstable side-to-side [3]. Most falls occur while walking [4, 5] and importantly, many of these falls occur during non-steady-state actions, such as incorrect weight shifting while attempting to maneuver [4]. In part, this high incidence of falls during maneuvers may result from the high frequency of such maneuvers during real-world walking [6]. Additionally, it has been suggested that humans trade off stability for maneuverability when performing lateral maneuvers [7, 8]. Humans enact walking maneuvers [9] and avoid falls [10] primarily by modulating their foot placement. Therefore, it is important to better understand how humans adapt stepping to simultaneously maintain balance and perform such maneuvers.

We developed a theoretical framework that defines repetitive tasks and proposes how humans regulate trial-to-trial variability with respect to Goal Equivalent Manifolds (GEMs) [11]. We used this framework to identify how humans regulate consecutive stepping movements during both straight-ahead walking [12, 13] and lateral maneuvers [14, 15]. In the lateral direction during straight-ahead walking, humans multi-objectively seek to minimize errors at each step with respect to primarily step width (*w*) to maintain lateral balance, as well as lateral body position (*z*_*B*_) to remain on their desired path [13]. However, if these task goals change substantially during lateral maneuvers, humans may need to abruptly and precisely modulate stepping in response. To enact a lateral lane-change maneuver, young healthy adults adapt stepping regulation at each step to directly trade off *w*-regulation for *z*_*B*_-regulation [14]. These results both support and explicitly quantify the proposed “stability-maneuverability trade-off” during lateral maneuvers [7, 8].

The limited studies that have assessed stepping strategies during walking maneuvers were primarily conducted with young healthy participants [7-9, 14]. However, older adults rely more on modulating lateral stepping to maintain lateral balance [16, 17]. Healthy aging is also associated with declines in both actual and perceived physical ability [5, 18], which in turn influence lateral stepping [19, 20]. Thus, while we previously found that young and older adults similarly regulate stepping during continuous walking [21], older adults may be less willing to enact the stability-maneuverability trade-offs exhibited by young adults during lateral maneuvers. Accordingly, the present analysis aimed to determine how older adults adapt stepping to enact lateral lane-change maneuvers, and to determine the influence of physical and perceived ability on their task performance.

## 2. Methods

### 2.1 Participants

Twenty young healthy (YH: 9M/11F, 21.7 ± 2.6 yrs) and eighteen older healthy (OH: 5M/13F, 71.6 ± 6.0 yrs) adults who participated in a previous experiment [22] were included in this analysis. All participants signed informed consent statements approved by the Institutional Review Board at The Pennsylvania State University prior to participating. All participants were screened to ensure they had no history of orthopedic problems, recent lower extremity injuries, visible gait anomalies, or medication regimens that may have influenced their gait. Groups did not differ in height, body mass, BMI, or leg length (all p > 0.25) [22].

### 2.2 Physical Ability and Perceived Ability Assessments

We assessed [22] participants’ balance and lower extremity strength using 30-Second Chair Stand (30CST) [23], Timed Up and Go (TUG) [24], and Four Square Step Test (FSST) [25] (Table 1). We assessed participants’ perceived balance ability using Activities-Specific Balance Confidence Scale (ABC) [26] and abbreviated Iconographic Falls Efficacy Scale (I-FES) [27]. For each grouping, we then calculated composite “Physical Ability” and “Perceived Ability” scores using principal component analysis (Table 1) [22, 28].

**Table 1:** Young healthy (YH) and older healthy (OH) participant assessment scores and composite Physical Ability and Perceived Ability scores [22]. Composite scores were computed using principal components analysis [28], where each composite score was defined as the linear combination of individual assessment z-scores that explained the most between-subject variance. All values are given as Mean ± Standard Deviation. Group mean differences were determined using two-sample t-tests or Mann-Whitney U tests [22]. Bold p-values indicate statistically significant differences.

| Characteristic: | Young Healthy (YH) | Older Healthy (OH) | p-value |
| --- | --- | --- | --- |
| **Physical Ability:** |  |  |  |
| 30CST [stands/30s] | 17.8 ± 6.0 | 13.7 ± 3.3 | **0.002** |
| TUG [s] | 6.62 ± 0.75 | 8.20 ± 1.9 | **0.004** |
| FSST [s] | 7.63 ± 1.4 | 9.19 ± 1.5 | **0.002** |
| Physical Composite =<br>0.496 * (30CST) – 0.636 * (TUG) – 0.592 * (FSST) | 0.1 ± 3.2 | -3.8 ± 3.2 | **< 0.001** |
| **Perceived Ability:** |  |  |  |
| ABC [%] | 96.8 ± 2.5 | 93.1 ± 5.5 | **0.023** |
| I-FES [/40] | 12.3 ± 1.9 | 16.1 ± 3.9 | **0.001** |
| Perceived Composite =<br>0.707 * (ABC) – 0.707 * (I-FES) | 2.3 ± 1.5 | -0.8 ± 3.2 | **< 0.001** |

### 2.3 Lateral Lane-Change Maneuver Task

Participants walked in an “M-Gait” system, comprised of a 1.2m wide motorized treadmill in a virtual reality environment (Motek, Amsterdam, Netherlands). Each participant walked at a constant speed of 0.75 m/s. Following a 4-minute acclimation trial, participants completed several different walking trials involving path navigation [22]. The data analyzed here were from one such trial, where participants were instructed to switch between two parallel paths, centered 0.6m apart, following an audible and visual cue (Fig. 1). Participants completed 6 such maneuvers in one 4-minute walking trial. They were instructed to walk normally on their current path between each maneuver. To ensure consistent task performance, we analyzed only the middle four maneuvers from each participant. Thus, we analyzed 80 lateral maneuvers from YH participants and 72 from OH participants.

**Figure 1:**
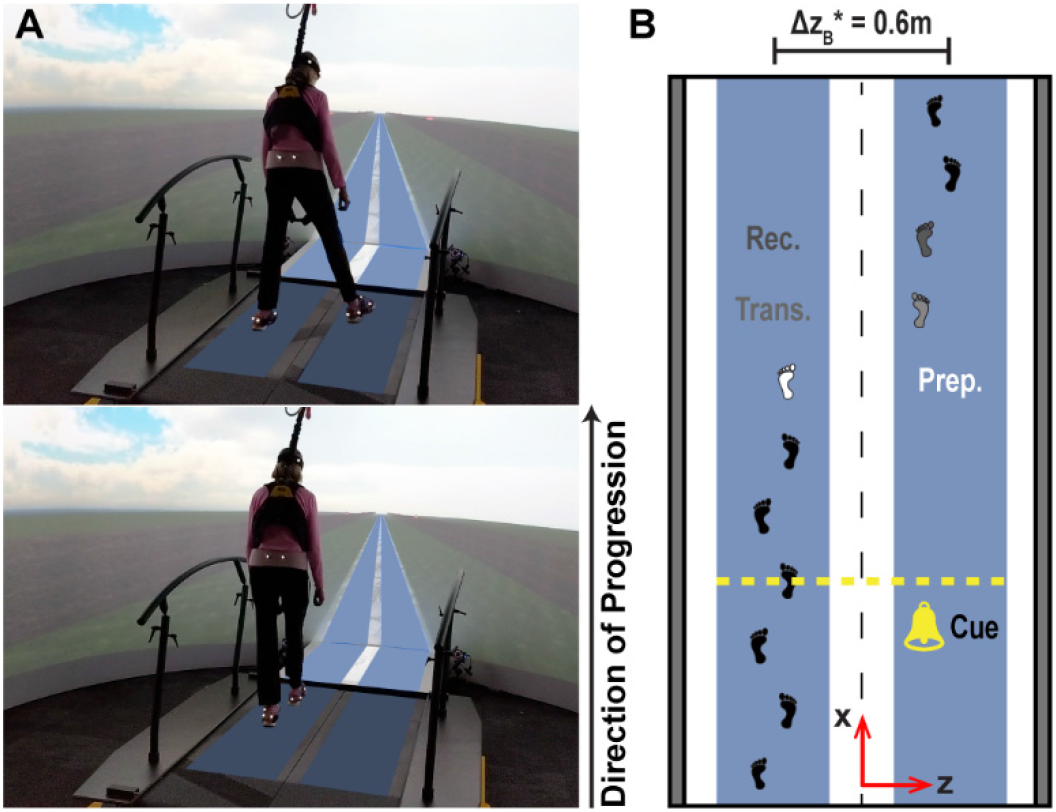
A) Participants were instructed to walk on one of two parallel paths projected onto the treadmill, the centers of which were 0.6m apart. They were instructed that, following an audible and visual cue, they were to switch (i.e., maneuver) to the adjacent path. B) A hypothetical example of a maneuver completed in four steps. Following two additional steps after the cue, the walker takes a small preparatory (Prep.) step, a large transition (Trans.) step to the new path, and a final recovery (Rec.) step before returning to straight-ahead walking on their new path. We previously demonstrated that young healthy adults typically utilize such a four-step strategy to perform lateral lane-change maneuvers [14].

### 2.4 Data Collection and Analyses

All relevant experimental walking data are available on Dryad [15]. Kinematic data were recorded from 16 retroreflective markers placed on the head, pelvis, and feet [22]. Marker trajectories were collected at 120Hz using a 10-camera Vicon motion capture system (Oxford Metrics, Oxford, UK) and post-processed using Vicon Nexus and D-Flow software (Motek, Amsterdam, Netherlands). Marker trajectories were analyzed in MATLAB (MathWorks, Natick, MA). Heel strikes were determined using a velocity-based detection algorithm [29]. Lateral foot placements (*z*_*L*_ and *z*_*R*_) were defined as the lateral location of the heel marker at each heel strike. Step width (*w*) was defined as the lateral displacement between foot placements: *w* = *z*_*R*_ − *z*_*L*_. Lateral position (*z*_*B*_) was defined as the midpoint between foot placements: *z*_*B*_ = 1/2(*z*_*L*_ + *z*_*R*_) [13].

### 2.5 Stepping Characteristics

We were unable to determine accurate heel strikes for three lateral maneuvers (one YH, two OH) and thus excluded these maneuvers from further analyses. One YH maneuver was completed with a different stepping strategy, a large cross-over step, and was therefore also excluded. Stepping data from the remaining 78 YH maneuvers and 70 OH maneuvers were normalized to a consistent direction (left-to-right).

To determine *when* participants performed these maneuvers, we computed the number of steps taken following the cue and before initiating the maneuver. We then aligned each maneuver to the initiation of the transition, defined here as the last step taken on the initial path. Mean *w* and *z*_*B*_ were determined for each step of the lateral maneuver.

Statistical analyses were performed using Minitab (Minitab, Inc, State College, PA). Group differences in the number of steps taken before initiating the maneuver were assessed using single-factor Mixed Effects Analyses of Variance (ANOVA). Group differences in mean *w* and *z*_*B*_ were assessed using two-factor (Group x Step) Mixed Effects ANOVAs. Individual Group and Step differences in mean *w* and *z*_*B*_ were determined using Tukey’s least significant difference pairwise comparisons. For each ANOVA, we then also added Physical Ability and Perceived Ability composite scores as covariates to determine these factors’ influence.

To determine *how* participants performed these maneuvers, we first plotted all lateral foot placements for each group in the [*z*_*L*_, *z*_*R*_] plane. We analyzed such plots for straight-ahead walking periods both before and after each maneuver and also for each step of the lateral maneuver [14]. At each of these steps, data across all subjects in each group were pooled to compute a single 95% prediction ellipse from these [*z*_*L*_, *z*_*R*_] data for each group at each step [14, 30]. We characterized each ellipse by its aspect ratio, area, and orientation angles (measured counterclockwise from the +*w* axis). The aspect ratio reflects the relative weighting of *w* and *z*_*B*_ regulation (for orientations ≈ 0°): larger aspect ratios indicate stronger prioritization of *w*-regulation. The area characterizes overall [*z*_*L*_, *z*_*R*_] stepping variability: larger areas reflect greater variability. The orientation reflects the degree of alignment to the *w* GEM: larger angles indicate less concern about minimizing deviations relative to the desired *w* [14]. We computed 95% confidence intervals for each ellipse characteristic from 10,000 bootstrapped samples, with replacement, for each group at each step.

As bootstrapping was required to obtain estimates of within-group variance for these ellipse characteristics, standard inferential statistics would not be appropriate. To compare these estimates between Groups, we instead used a previously established two-sample procedure for bootstrapped data [31]. The YH and OH stepping data were pooled, from which we generated 10,000 new bootstrapped samples with replacement at each step. These pooled, bootstrapped data were randomly split into two groups, sized in accordance with the experimental data. This generated a series of between-group differences with distribution ∼*N(0,S*_*P*_) for each ellipse characteristic at a given step. A p-value was then computed as the percentage of bootstrapped samples with a greater difference than that observed experimentally [31].

## 3. Results

For both groups, approximately half of the cues prompting participants to switch paths occurred on a contralateral step with respect to the transition direction, while the other half occurred on an ipsilateral step (Fig. 2A-B). Both YH and OH participants took more steps following cues presented on an ipsilateral step than a contralateral step (p < 0.001). The number of steps was not different between Groups when pooling all maneuvers (p = 0.076), nor when separating maneuvers by contralateral (p = 0.063) versus ipsilateral (p = 0.148) cue step. When we added both Physical Ability and Perceived Ability scores as covariates in these models, the number of steps was not influenced by Group (all p > 0.53) nor Physical Ability (all p > 0.45). However, *Perceived Ability* significantly influenced the number of steps taken following the cue when pooling across all maneuvers (p = 0.019) and for maneuvers cued on an ipsilateral step (p = 0.025), but not those cued on a contralateral step (p = 0.085). Participants with lower Perceived Ability scores tended to take more steps following the cue before initiating these maneuvers (Fig. 3A).

**Figure 2:**
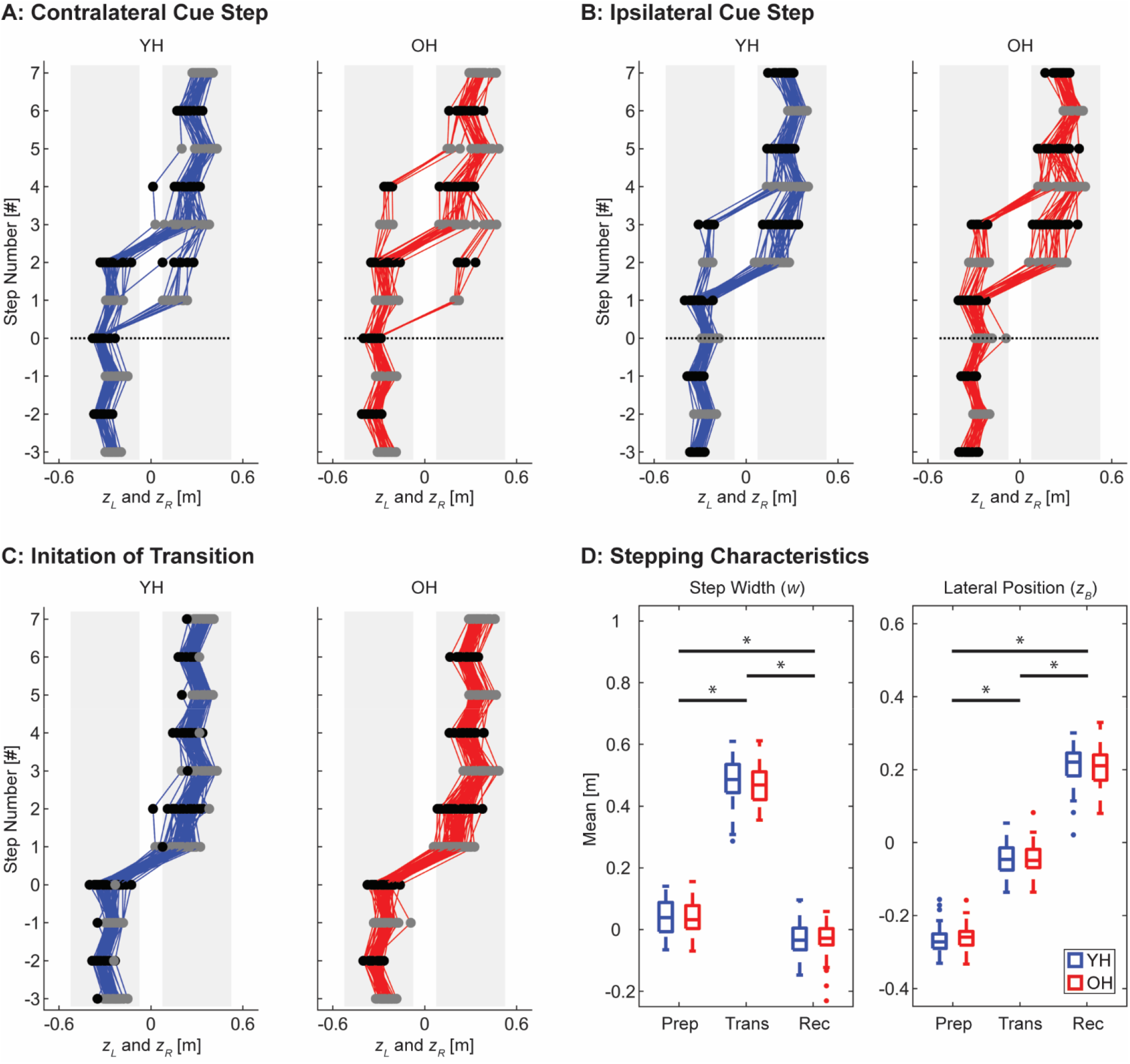
A) Time series of Young Healthy (YH) and Older Healthy (OH) left (*z*_*L*_; black) and right (*z*_*R*_; gray) foot placements for transitions cued on a contralateral step relative to the direction of transition [38 of 79 total transitions (48%) for YH; 33 of 70 (47%) for OH]. B) Analogous time series of foot placements for transitions cued on an ipsilateral step relative to the direction of transition [41 of 79 (52%) for YH; 37 of 70 (53%) for OH]. For both (A) and (B), the black dotted lines indicate the onset of the cue to transition. C) Time series of all transitions for both YH and OH plotted with respect to when they initiated the transition, defined as the last step taken on the original path (step 0). All transitions are plotted to appear from left to right. D) Step widths (*w*) and lateral body positions (*z*_*B*_) at the preparatory (0), transition (1), and recovery (2) steps for YH and OH participants. The summary boxplots indicate the median, 1^st^ and 3^rd^ quartiles, and whiskers extending to 1.5 x interquartile range. Values beyond this range are shown as individual data points. For both *w* and *z*_*B*_, differences between Steps were highly significant (both p < 0.001). However, Group differences were not (both p > 0.44).

**Figure 3:**
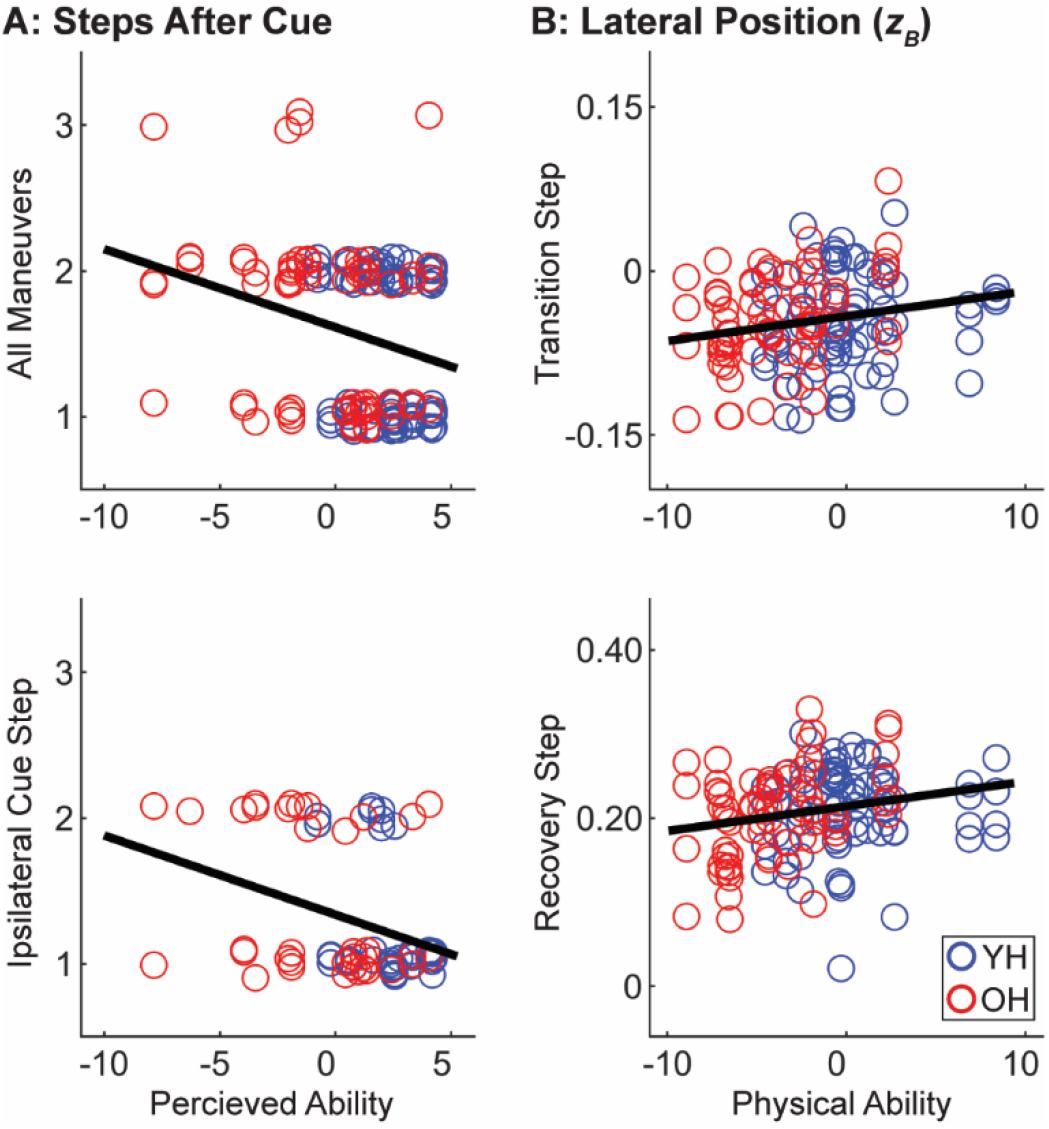
A) Correlations between Perceived Ability composite scores and the number of contralateral steps taken following the cue for all maneuvers (top) and those cued on an ipsilateral step (bottom). B) Correlations between Physical Ability composite scores and lateral position (*z*_*B*_) at the transition (top) and recovery (bottom) steps of the lateral maneuver. Least squares regression lines shown on each plot are for reference only to indicate the direction of each relationship.

Consistent with our prior results [14], both YH and OH participants performed nearly all maneuvers consistently once initiated (Fig. 2C). Except for the one YH participant who completed one maneuver with a large cross-over step, all other maneuvers were completed with a four-step strategy. Participants first took an initial “preparatory” step (step 0), where they narrowed their step width and incrementally moved towards their new path. Participants took a large “transition” step (step 1) onto their new path. Participants took a “recovery” step (step 2) that again exhibited a slightly narrower step width. Participants then reached their new final position and retuned to straight-ahead walking (step 3).

Both mean step width and mean lateral position significantly differed at each of the preparatory, transition, and recovery steps (both p < 0.001). However, Group differences were not significant for either variable (both p > 0.44; Fig. 2D). When composite ability scores were included as covariates in these models, step width was not significantly influenced by Physical Ability (p = 0.605) nor Perceived Ability (p = 0.128). However, *Physical Ability* significantly influenced lateral position (p = 0.031), but Perceived Ability did not (p = 0.336). Participants with greater Physical Ability scores tended to have a greater lateral position at the transition and recovery steps, indicating that they reached their new path faster (Fig. 3B).

The OH ellipses were significantly more isotropic than the YH ellipses during straight-ahead walking both before (p = 0.035) and after (p = 0.001) the lateral maneuvers (Fig. 4C). However, the ellipses of both groups were strongly aligned to the constant step width (*w*^*\**^) GEM (Fig. 4A), consistent with our prior work demonstrating a strong prioritization for *w*-regulation [13]. No significant Group differences were observed before or after the maneuver in ellipse areas (p = 0.211 and p = 0.242) or orientations (p = 0.311 and p = 0.052).

**Figure 4:**
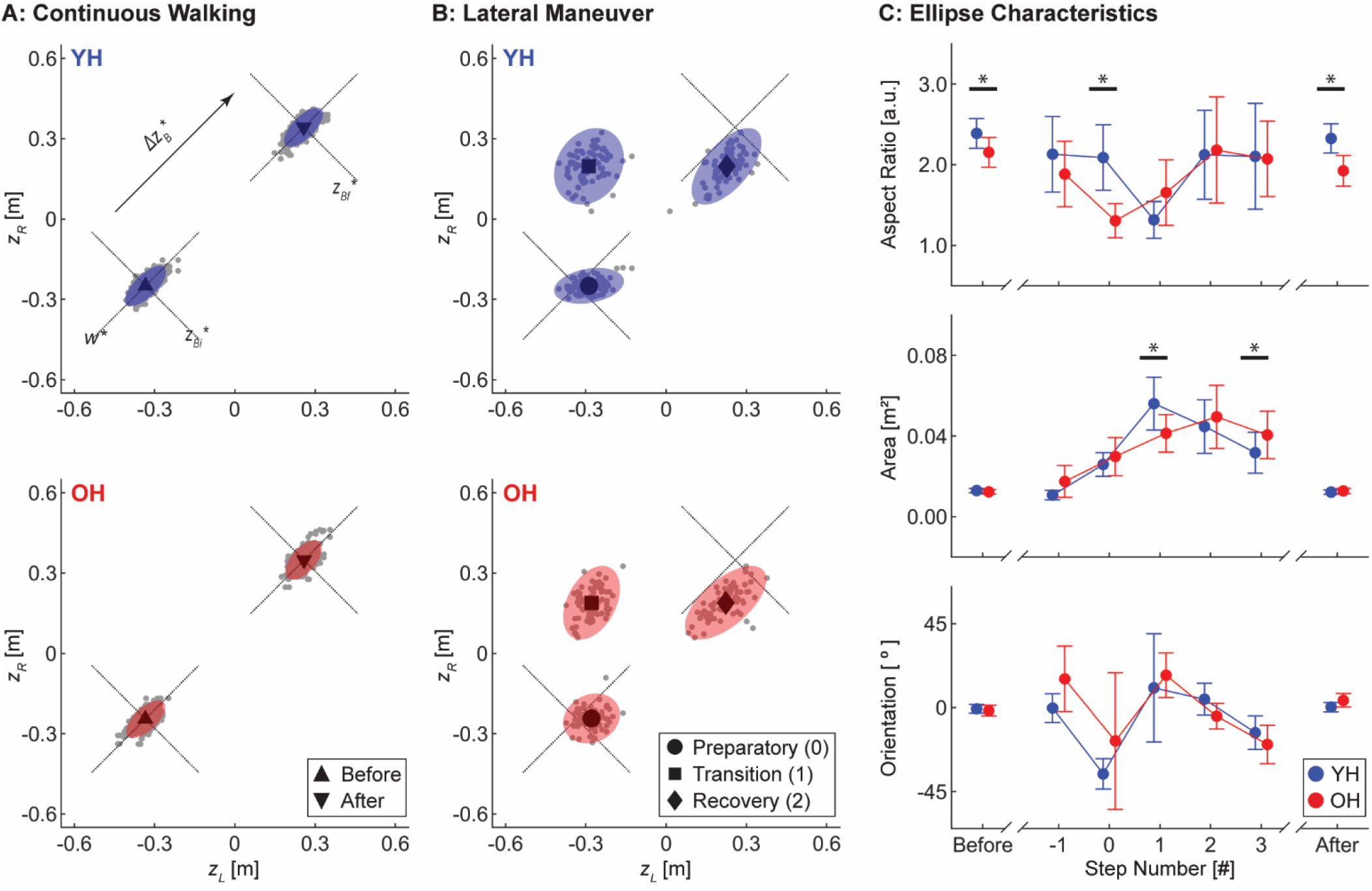
A) Stepping data from Young Healthy (YH) and Older Healthy (OH) participants during the straight walking periods before and after the transition, projected onto the [*z*_*L*_, *z*_*R*_] plane. For each group, data were pooled across 8 steps before and 8 steps after each of 4 transitions from each participant. B) Stepping data during the preparatory, transition, and recovery steps from 78 analyzed YH maneuvers and 70 analyzed OH maneuvers projected onto the [*z*_*L*_, *z*_*R*_] plane, plotted to appear from left to right. For both (A) and (B), the diagonal dotted lines indicate the initial (*z*_*Bi*_^*\**^) and final (*z*_*Bf*_^*\**^) constant-*z*_*B*_^*^ (downward sloping) and constant-*w*^*^ (upward sloping) GEMs, determined from averages of each group. Gray dots indicate individual steps, and the blue (YH) and red (OH) ellipses depict 95% prediction ellipses. C) Prediction ellipse characteristics during straight-ahead walking and at each step of the lane-change maneuver for YH and OH participants: aspect ratio (top), area (center), and orientation (bottom). Error bars indicate ±95% confidence intervals at each step derived from 10,000 bootstrapped samples. Statistically significant group differences (p < 0.05) [31] are indicated by the asterisks (*).

During the lateral maneuver itself, OH participants exhibited a significantly lower ellipse aspect ratio at the preparatory step (p < 0.001), demonstrating that they began trading off *w*-regulation for *z*_*B*_-regulation prior to initiating the maneuver (step 0; Fig. 4B-C). Ellipse aspect ratios did not differ at the subsequent transition step (step 1; p = 0.211), as the OH ellipse remained more isotropic and the YH ellipse demonstrated a similar reduction in aspect ratio (Fig. 4B-C). Thus, both groups similarly traded-off *w*-for *z*_*B*_-regulation to enact the lateral lane-change. The area of the YH ellipse was significantly greater than that of the OH ellipse at the transition step (p = 0.044), indicating greater overall stepping variability among YH participants. However, YH participants reduced stepping variability faster than OH participants following the maneuver, evident by a significantly greater area of the OH ellipse at step 3 (p = 0.047; Fig. 4C).

## 4. Discussion

Humans frequently perform maneuvers when walking in the real world [6]. Prior work suggested humans trade off stability to enact such maneuvers [7, 8], potentially contributing falls among older adults during these types of tasks [4]. Humans perform maneuvers [9] and avoid falls [10] primarily by modulating their foot placement. We previously developed a theoretical framework to determine how young adults adapt stepping from each step to the next during lateral lane-change maneuvers to accomplish both objectives (stability & maneuverability) simultaneously [14]. Here, we applied this framework to determine how healthy older adults perform this task.

Older adults typically used the same four-step strategy as young adults to perform the lateral maneuver (Fig. 2). No significant Group differences were observed in either step width or lateral position at the preparatory, transition, or recovery steps, nor did Perceived Ability influence these measures. However, lateral positions at the transition and recovery steps were significantly greater among participants with greater *Physical* Ability scores (Fig. 3B), corroborating our prior work demonstrating the influence of physical ability on walking task performance [22]. Given the substantial impulses required to execute the lateral maneuver, it is unsurprising that participants with greater strength and mobility reached their new path more quickly. Nevertheless, these results demonstrate that physical ability, more than age *per se*, influenced how participants enacted the lateral lane-change.

The stepping ellipses were generally consistent between YH and OH participants, indicating similar stepping strategies among both groups. This result extends previous analogous findings for continuous walking [21]. During continuous walking both before and after the maneuver, however, the OH ellipse was slightly but significantly more isotropic than the YH ellipse (Fig. 4C). This result was surprising, as prior work has demonstrated a greater reliance on stepping to maintain lateral balance among older adults [16, 17]. However, the OH ellipses during continuous walking remained highly anisotropic and strongly aligned to the *w*^*^ GEM relative to those at the preparatory and transition steps (Fig. 4A-B), indicating that older adults still strongly prioritized *w*-regulation during continuous walking. Older adults typically exhibit deficits in locomotor planning, including prioritizing future over current stepping performance during complex walking tasks [32]. Thus, the difference in isotropy observed here may suggest that OH participants slightly adapted their stepping throughout the trial in anticipation of the frequent maneuvers.

The OH stepping ellipse became significantly more isotropic at the preparatory step prior to initiating the maneuver (Fig. 4C). This is consistent with the idea that these OH adults likely prioritized planning for the upcoming maneuver over current stepping performance [32]. Conversely, young adults did not substantially deviate from their continuous stepping regulation strategy until the transition step (Fig. 4C), indicating that they were able to plan for the maneuver and maintain *w*-regulation simultaneously. Nevertheless, the ellipses of both older and young adults were more isotropic at the transition step than during straight-ahead walking, indicating that both groups traded off *w*-regulation for *z*_*B*_-regulation to enact the maneuver. These results quantitatively capture the stability-maneuverability trade-off previously proposed to occur during lateral maneuvers [7, 8] and, to our knowledge, are the first to demonstrate how this trade-off differs for older adults.

The task assessed here required participants to not only adapt their stepping to enact the lateral lane-change, but also to decide when to initiate the maneuver. The varying number of steps taken following the cue before transitioning to the new path (Fig. 2) suggests that this decision likely varied across participants. While older adults took slightly more steps prior to initiating the maneuver, this difference did not reach statistical significance. However, lower *Perceived* Ability (across both groups) significantly predicted the increased number of steps taken prior to initiating the maneuver (Fig. 3A). Indeed, perception of one’s physical ability is a critical element of “embodied decisions” [33] and has been shown to influence decision-making among older adults during postural tasks [34], mobility-related tasks [18], and our prior work assessing a similar lateral maneuver task [22]. The present results extend these findings and suggest that participants’ intrinsic perception of their ability, more than age itself or actual physical ability, influenced *when* participants chose to initiate the maneuver.

Goal-directed walking, including the lateral maneuver assessed here, requires humans to make decisions related to the task of walking itself. These decisions are based not only on physical ability, but also perceptual judgments and executive function [33]. Here, the increased isotropy of the OH ellipses during both continuous walking and at the preparatory step prior to the maneuver (Fig. 4), as well as the varying number of steps taken following the cue before transitioning to the new path (Fig. 2), suggest that planning and decision-making during this task likely differed across participants. Furthermore, this work demonstrates that physical and perceived ability, both of which decline with healthy aging [5, 18], significantly influenced how humans perform lateral maneuvers, but in different ways. Perceived ability more strongly influenced *when* participants maneuvered, whereas physical ability more strongly influenced *how* they did so. Thus, the present results yield a better understanding of these interactions and may help to inform interventions to improve walking performance among older adults.

## Acknowledgments

The authors thank Ms. Anna C. Render for her assistance with data collection. This project was funded by the US National Institutes of Health / National Institute on Aging (NIA; Grant # R01-AG049735 & R21-AG053470; to JPC and JBD).

## References

[1] E. Burns, R. Kakara, Deaths from Falls Among Adults ≥65 Years—United States, 2007–2016, MMWR Morb. Mortal Weekly Report 67 (2018) 504–519. http://dx.doi.org/10.15585/mmwr.mm6718a1

[2] J.L. Kelsey, W.S. Browner, D.G. Seeley, M.C. Nevitt, S.R. Cummings, Risk factors for fractures of the distal forearm and proximal humerus. The study of osteoporotic fractures research group, Am J Epidemiol 135(5) (1992) 477–489. https://doi.org/10.1093/oxfordjournals.aje.a116314

[3] A.D. Kuo, Stabilization of lateral motion in passive dynamic walking, Int. J. Robotics Res. 18(9) (1999) 917–930. https://doi.org/10.1177/02783649922066655

[4] S.N. Robinovitch, F. Feldman, Y. Yang, R. Schonnop, P.M. Leung, T. Sarraf, et al., Video capture of the circumstances of falls in elderly people residing in long-term care: an observational study, Lancet 381(9860) (2013) 47–54. https://doi.org/10.1016/S0140-6736(12)61263-X

[5] J.L. Kelsey, E. Procter-Gray, M.T. Hannan, W. Li, Heterogeneity of Falls Among Older Adults: Implications for Public Health Prevention, Am. J. Public Health 102(11) (2012) 2149–2156. https://doi.org/10.2105/ajph.2012.300677

[6] B.C. Glaister, G.C. Bernatz, G.K. Klute, M.S. Orendurff, Video task analysis of turning during activities of daily living, Gait Posture 25(2) (2007) 289–94. https://doi.org/10.1016/j.gaitpost.2006.04.003

[7] J. Acasio, M.M. Wu, N.P. Fey, K.E. Gordon, Stability-maneuverability trade-offs during lateral steps, Gait Posture 52 (2017) 171–177. https://doi.org/10.1016/j.gaitpost.2016.11.034

[8] M. Wu, J.H. Matsubara, K.E. Gordon, General and specific strategies used to facilitate locomotor maneuvers, PLoS ONE 10(7) (2015) e0132707. https://doi.org/10.1371/journal.pone.0132707

[9] K.L. Hsieh, R.C. Sheehan, J.M. Wilken, J.B. Dingwell, Healthy individuals are more maneuverable when walking slower while navigating a virtual obstacle course, Gait Posture 61 (2018) 466–472. https://doi.org/10.1016/j.gaitpost.2018.02.015

[10] S.M. Bruijn, J.H. van Dieën, Control of human gait stability through foot placement, J. R. Soc. Interface 15(143) (2018) 1–11. https://doi.org/10.1098/rsif.2017.0816

[11] J.P. Cusumano, J.B. Dingwell, Movement variability near goal equivalent manifolds: Fluctuations, control, and model-based analysis, Hum. Mov. Sci. 32(5) (2013) 899–923. https://doi.org/10.1016/j.humov.2013.07.019

[12] J.B. Dingwell, J. John, J.P. Cusumano, Do humans optimally exploit redundancy to control step variability in walking?, PLoS Comput. Biol. 6(7) (2010) e1000856. https://doi.org/10.1371/journal.pcbi.1000856

[13] J.B. Dingwell, J.P. Cusumano, Humans use multi-objective control to regulate lateral foot placement when walking, PLoS Comput. Biol. 15(3) (2019) e1006850. https://doi.org/10.1371/journal.pcbi.1006850

[14] D.M. Desmet, J.P. Cusumano, J.B. Dingwell, Adaptive multi-objective control explains how humans make lateral maneuvers while walking, PLoS Comput Biol 18(11) (2022) e1010035. https://doi.org/10.1371/journal.pcbi.1010035

[15] [dataset] [15] D.M. Desmet, J.P. Cusumano, J.B. Dingwell, Data from: Adaptive multi-objective control explains how humans make lateral maneuvers while walking, Dryad, Dataset, 2022. https://doi.org/10.5061/dryad.tx95×6b1x

[16] A. Vistamehr, R.R. Neptune, Differences in balance control between healthy younger and older adults during steady-state walking, J. Biomech. 128 (2021) 110717. https://doi.org/10.1016/j.jbiomech.2021.110717

[17] T.M. Owings, M.D. Grabiner, Variability of step kinematics in young and older adults, Gait Posture 20(1) (2004) 26–29. https://doi.org/10.1016/s0966-6362(03)00088-2

[18] B.L. Fischer, C.E. Gleason, R.E. Gangnon, J. Janczewski, T. Shea, J.E. Mahoney, Declining Cognition and Falls: Role of Risky Performance of Everyday Mobility Activities, Phys. Ther. 94(3) (2014) 355–362. https://doi.org/10.2522/ptj.20130195

[19] B.E. Maki, Gait Changes in Older Adults: Predictors of Falls or Indicators of Fear?, Journal of the American Geriatric Society 45(3) (1997) 313–320. https://doi.org/10.1111/j.1532-5415.1997.tb00946.x

[20] S. Shin, J. Valentine, E. Evans, J. Sosnoff, Lower extremity muscle quality and gait variability in older adults, Age and Ageing 41 (2012) 595–599. https://doi.org/10.1093/ageing/afs032

[21] M.E. Kazanski, J.B. Dingwell, J.P. Cusumano, How healthy older adults regulate lateral foot placement while walking in laterally destabilizing environments, J. Biomech. 104 (2020) 109714. https://doi.org/10.1016/j.jbiomech.2020.109714

[22] M.E. Kazanski, J.B. Dingwell, Effects of age, physical and self-perceived balance abilities on lateral stepping adjustments during competing lateral balance tasks, Gait Posture 88 (2021) 311–317. https://doi.org/10.1016/j.gaitpost.2021.05.025

[23] C. Jones, R. Rikli, W. Beam, A 30-s chair-stand test as a measure of lower body strength in community-residing older adults, Res Q Exerc Sport 70(2) (1999) 113–119. https://doi.org/10.1080/02701367.1999.10608028

[24] D. Podsiadlo, S. Richardson, The Timed Up and Go Test - A Test of Basic Functional Mobility for Frail Elderly Persons, J. Am. Geriatr. Soc. 39(2) (1991) 142–148. https://doi.org/10.1111/j.1532-5415.1991.tb01616.x

[25] W. Dite, V.A. Temple, A clinical test of stepping and change of direction to identify multiple falling older adults, Arch. Phys. Med. Rehabil. 83(11) (2002) 1566–1571. http://dx.doi.org/10.1053/apmr.2002.35469

[26] L.E. Powell, A.M. Myers, The Activities-specific Balance Confidence (ABC) Scale, The Journals of Gerontology Series A: Biological Sciences and Medical Sciences 50A(1) (1995) M28–M34. https://doi.org/10.1093/gerona/50A.1.M28

[27] K. Delbaere, S. T. Smith, S.R. Lord, Development and Initial Validation of the Iconographical Falls Efficacy Scale, The Journals of Gerontology Series A: Biological Sciences and Medical Sciences 66A(6) (2011) 674–680. https://doi.org/10.1093/gerona/glr019

[28] H.G. Kang, J.B. Dingwell, Separating the effects of age and speed on gait variability during treadmill walking, Gait Posture 27(4) (2008) 572–577. https://doi.org/10.1016/j.gaitpost.2007.07.009

[29] J.A. Zeni, J.G. Richards, J.S. Higginson, Two simple methods for determining gait events during treadmill and overground walking using kinematic data, Gait Posture 27(4) (2008) 710–714. https://doi.org/10.1016/j.gaitpost.2007.07.007

[30] P. Schubert, M. Kirchner, Ellipse area calculations and their applicability in posturography, Gait Posture 39(1) (2014) 518–22. https://doi.org/10.1016/j.gaitpost.2013.09.001

[31] T. Hesterberg, Bootstrap, WIREs Computational Statistics 3(6) (2011) 497–526. https://doi.org/10.1002/wics.182

[32] G.J. Chapman, M.A. Hollands, Evidence that older adult fallers prioritise the planning of future stepping actions over the accurate execution of ongoing steps during complex locomotor tasks, Gait Posture 26(1) (2007) 59–67. http://dx.doi.org/10.1016/j.gaitpost.2006.07.010

[33] J. Gordon, A. Maselli, G.L. Lancia, T. Thiery, P. Cisek, G. Pezzulo, The road towards understanding embodied decisions, Neuroscience & Biobehavioral Reviews 131 (2021) 722–736. https://doi.org/10.1016/j.neubiorev.2021.09.034

[34] G. Lafargue, M. Noel, M. Luyat, In the elderly, failure to update internal models leads to over-optimistic predictions about upcoming actions, PLoS ONE 8(1) (2013) e51218. https://doi.org/10.1371/journal.pone.0051218

